# Interpretable deep recommender system model for prediction of kinase inhibitor efficacy across cancer cell lines

**DOI:** 10.1101/2021.01.26.428272

**Authors:** Krzysztof Koras, Ewa Kizling, Dilafruz Juraeva, Eike Staub, Ewa Szczurek

## Abstract

Computational models for drug sensitivity prediction have the potential to revolutionise personalized cancer medicine. Drug sensitivity assays, as well as profiling of cancer cell lines and drugs becomes increasingly available for training such models. Machine learning methods for drug sensitivity prediction must be optimized for: (i) leveraging the wealth of information about both cancer cell lines and drugs, (ii) predictive performance and (iii) interpretability. Multiple methods were proposed for predicting drug sensitivity from cancer cell line features, some in a multi-task fashion. So far, no such model leveraged drug inhibition profiles. Recent neural network-based recommender systems arise as models capable of predicting cancer cell line response to drugs from their biological features with high prediction accuracy. These models, however, require a tailored approach to model interpretability. In this work, we develop a neural network recommender system for kinase inhibitor sensitivity prediction called DEERS. The model utilizes molecular features of the cancer cell lines and kinase inhibition profiles of the drugs. DEERS incorporates two autoencoders to project cell line and drug features into 10-dimensional hidden representations and a feed-forward neural network to combine them into response prediction. We propose a novel model interpretability approach offering the widest possible assessment of the specific genes and biological processes that underlie the action of the drugs on the cell lines. The approach considers also such genes and processes that were not included in the set of modeled features. Our approach outperforms simpler matrix factorization models, achieving R=0.82 correlation between true and predicted response for the unseen cell lines. Using the interpretability analysis, we evaluate correlation of all human genes with each of the hidden cell line dimensions. Subsequently, we identify 67 biological processes associated with these dimensions. Combined with drug response data, these associations point at the processes that drive the cell line sensitivity to particular compounds. Detailed case studies are shown for PHA-793887, XMD14-99 and Dabrafenib. Our framework provides an expressive, multitask neural network model with a custom interpretability approach for inferring underlying biological factors and explaining cancer cell response to drugs.

## Background

Matching the optimal drugs for individual cancer patients remains a crucial problem of precision medicine (1). Drug sensitivity data from cancer models are frequently generated to provide the basis for the discovery of molecular markers to predict drug efficacy. To predict the response of a specific cell line to a specific drug, there is a need of computational models that can leverage the abundance of information about drugs and cancer cell lines.

Strongly parallelized assay formats provide a variety of data that can be used to comprehensively describe the characteristics of both cancer cell lines and drugs (2–6). Despite their drawbacks (7–11), multi-omics cell line data can provide important insights about the molecular mechanisms underlying susceptibility to distinct drugs. Arguably, the subclass of kinase inhibitor drugs is best characterized by their kinase inhibition profiles, which, apart from the intended on-targets, manifest also off-target effects. Despite their frequent use during the early phases of drug development, when inhibitory profiles of kinase inhibitors are optimized, to our knowledge such data has not been used for modelling of drug response. Computational drug sensitivity prediction has been approached by many machine learning methodologies (12–14), ranging from traditional algorithms (15–18) to models based on neural networks and deep learning (19–24).

The problem of drug sensitivity prediction can be stated as a recommendation problem, where cancer cell lines and drugs are analogous to users and items, respectively. The goal is to recommend the best drug for a given cell line. One of the most popular recommender system techniques is matrix factorization (MF), where the user-item interaction matrix is decomposed into a product of two lower-dimensional rectangular matrices. The problem of so called matrix factorization with side information incorporates features of users and items in the factorization process. The simplest approach to such MF problems involves linear projection of the features to lower-dimensional hidden space, followed by computing the dot product between corresponding user and item hidden representations in order to obtain user-item interaction prediction (25–27). Recently, this approach has been modified by introducing non-linearity in the projection step, where the projections are computed by neural networks or autoencoders, but the corresponding hidden representations are still connected via a dot product in the linear fashion. Dot product, however, as a simple linear function, has a limited ability to capture the complex user-item interactions in the hidden space. To address this issue, deep neural networks have been proposed to replace the dot product for modeling the user-item interactions in the latent space (28, 29). Since neural networks are known as the universal approximators (30), they are expected to be more suitable to learn complex relationships between the hidden representations of the users and items and the response variable.

While the neural-network based models are more expressive, previous analyses point out that the deep learning models do not necessarily outperform simpler models when the latter are finely tuned, and that some published neural network model results are hard to reproduce (31). Moreover, deep neural networks have a reputation of being difficult to interpret due to their non-linearity and complex structure. The majority of so called explainable artificial intelligence methods focus on finding attributions between specific neurons in the network by analyzing the underlying gradient flow (32– 35). Although useful, these methods provide rather standard utilities (e.g. feature importances), often available also for traditional machine learning models. Moreover, the insights derived from such interpretability approaches are limited by the features chosen for training the model.

We argue that a desired recommender system for the problem of drug sensitivity prediction should satisfy several objectives. First, it should solve a multi-task learning problem, i.e. model multiple drugs and cell lines simultaneously. This allows to capture general mechanisms driving the drug-cell lines interactions. Second, it should achieve state-of-the art predictive performance, especially in the task of predicting drug sensitivities for new cell lines. This is due to the fact that in this setting, the new cell line mimics a new patient, and the recommendation problem corresponds to identifying the best therapy for that patient. Finally, the model should be interpretable. Specifically, the model should explain the rationale behind its predictions and provide biological and pharmacological insights regarding the mechanism underlying the drugs-cell lines interactions. The emphasis on model interpretability is crucial in the context of its potential clinical applications.

To address these objectives, we develop a recommender system model for drug sensitivity prediction, called DEERS (Drug Efficacy Estimation Recommender System). DEERS incorporates two autoencoders to project the drug and cell line features, respectively, into lower dimensional representations, and uses a feed forward network to predict the sensitivities of the cell lines to the drugs based on their hidden representations. The proposed framework brings several advantages. First, the model solves a multi-drug and multi-cell line sensitivity learning problem and utilizes cell lines biological data and drugs inhibition profiles as side information (Fig. 1a,b). Second, the model is highly predictive. In a comparative analysis, DEERS outperforms two other MF-based recommender system models, and achieves similarly good results to the best performing state-of-the art XGBoost algorithm. Third, we provide an approach for model interpretability, on two levels: i) meaningful drug and cell line feature representation learning, and ii) explaining the cell line sensitivities to drugs in terms of the underlying biological processes.

**Fig 1.**
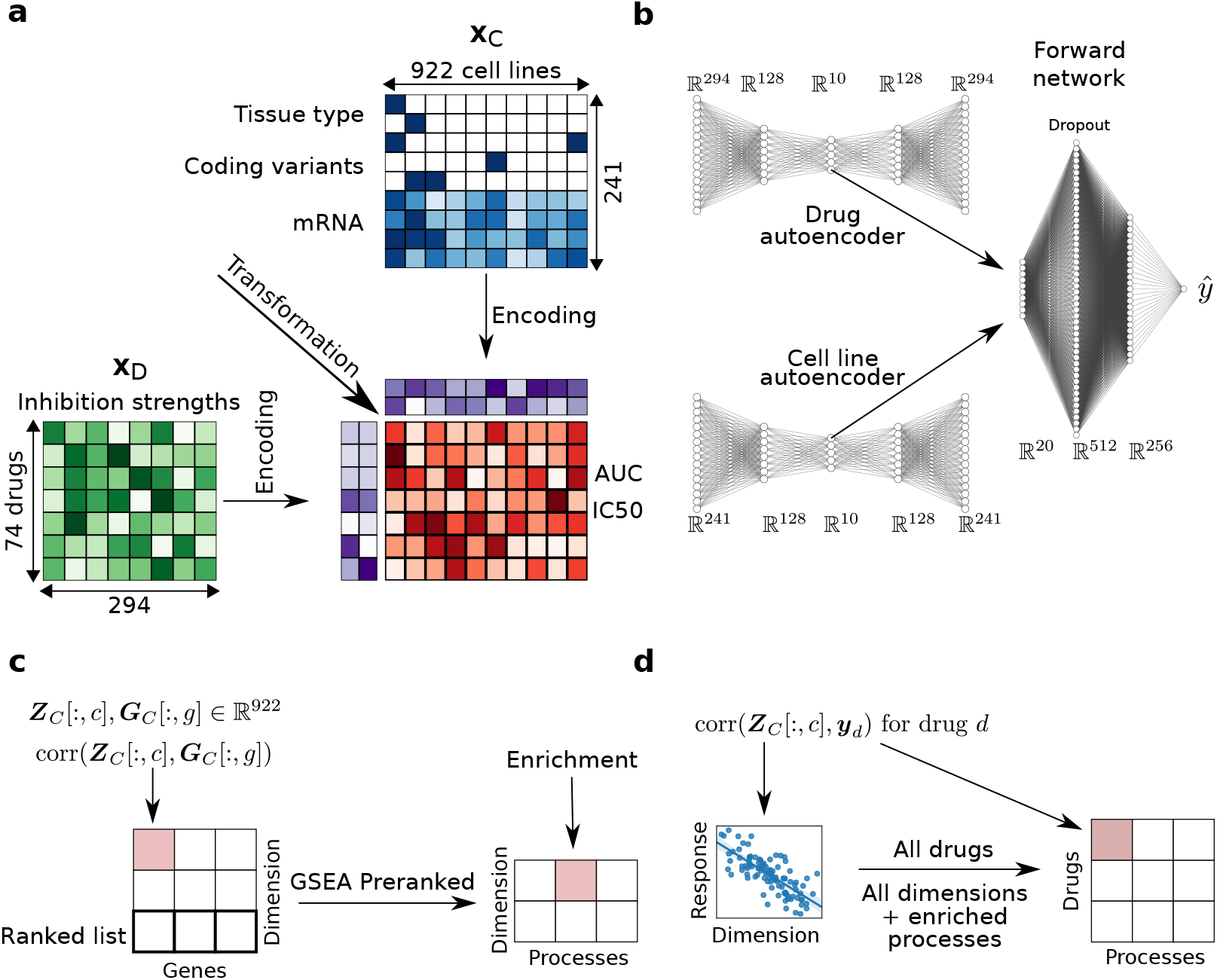
Overview of the data and the modeling process. (**a**) Recommender system framework for drug sensitivity prediction from drug and cell line features. The drugs are described by their inhibition profiles on a panel of 294 kinases. The biological features of the cell lines include continuous mRNA expressions, binary indicators of coding variants, and dummy-encoded tissue type. Two drug response metrics are considered: AUC and IC50. The recommender system first independently encodes drugs and cell lines input data into lower-dimensional representations. The two hidden representations are then transformed in order to compute the drug response estimation. (**b**) Architecture of the DEERS model. First, the drugs and cell lines inputs are passed into corresponding autoencoders which output the 10-dimensional representations and reconstructed data. Next, the hidden representations are concatenated and used as an input to the two-layered, feed-forward network which outputs the drug response estimate. (**c**) Method for relating biological meaning to hidden dimensions of cell lines. First, the hidden dimensions of the cell line autoencoder are correlated with gene expression data. Here, ***Z***_*C*_ denotes the matrix with cell line hidden representations stacked in rows, ***Z***_*c*_[:, *c*] denotes a column of ***Z***_*C*_, ***G***_*C*_ denotes the gene expression data for cell lines and ***G***_*C*_ [:, *g*] denotes a column of ***G***_*C*_. The resulting ranked lists, one per each dimension, are then passed as an input to GSEA Preranked analysis, obtaining biological processes enriched in every hidden dimension (see Methods). (**d**) Method for relating the drug action directly to biological processes. For a given drug *d*, the cell line response is correlated with a given cell line hidden dimension *c*. The obtained correlation coefficient is then mapped to the biological processes enriched in hidden dimension *c*. This procedure is performed for every drug and every hidden dimension, obtaining the matrix relating drugs to the biological processes (see Methods).

The crucial aspect of the proposed interpretability approach is that it offers the widest possible assessment of the specific genes and biological processes that underlie the action of the drugs on the cell lines. The novelty of this approach stems from the fact that it considers also such genes and processes that were not included in the set of modeled features. Using the interpretablity approach, we demonstrate that the low-dimensional representations of the model capture the high dimensional features of drugs or cell lines, specifically the molecular patterns of cell lines and drug inhibition profiles that govern the response of distinct cell lines to drugs (Fig. 1c). Finally, we find the relationships between drug response and biological processes of cell lines (Fig. 1d).

## Methods

### Analyzed data

The analyzed dataset comprised measurements of drug sensitivity of cell lines using viability assays for a total of 922 cell lines and 74 drugs, corresponding to 52,730 drug-cell line pairs. The sensitivity measurements were acquired from the Genomics of Drug Sensitivity in Cancer (GDSC) (3) database. GDSC provides two sensitivity measurements, summarizing the dose-response curve: area under the curve (AUC) and log half maximal inhibitory concentration (IC50), defined as a drug concentration needed to reduce cell viability by 50%. Both sensitivity metrics were used to train and assess the performance of the presented models. Drug sensitivity of a cell line is the prediction target of our modeling approach.

The group of 74 drugs selected for modeling consisted exclusively of kinase inhibitors. The drugs in this group differ from other cancer drugs by their mode of action. Data to characterize the 74 kinase inhibitors were extracted from the HMS LINCS KINOMEscan data resource (36). The features set of these drugs consisted of binding strength across a panel 294 protein kinases (Fig. 1a). The value for a given compound-kinase pair represents a percent of control, where a 100% result means no inhibition of kinase binding to the ligand in the presence of the compound, and where low percent results mean strong inhibition (37, 38). The data was acquired for those 74 drugs which were also present in the GDSC database, yielding a final drug characterization matrix for 74 drugs and 294 protein kinases.

Data to characterize the 922 cell lines were downloaded from the GDSC. For the molecular features of the cell lines, we considered only the genes coding for kinases present in KI-NOMEscan dataset, as well as any putative gene targets of all considered compounds. This resulted in the set of 202 genes, for which mRNA expression levels (202 features) and binary mutation calls (21 features) were extracted for all cell lines. Furthermore, the dummy-encoded tissue type was added, producing additional 18 binary features, yielding the final set of 241 biological features for 922 cell lines (Fig. 1a).

### DEERS: a deep neural network model of drug sensitivity accounting for inhibition of protein kinases by drugs and cancer cell line features

The goal of the proposed model is to predict a response of a given cell line to a given drug, i.e. estimate the corresponding AUC or IC50 value, given the drug and cell line feature representations (Fig. 1a). The final prediction is computed in two steps: first, we compute lower-dimensional representations of the considered drug and cell line, and second, the representations are combined, in order to make the sensitivity estimation. This problem can be viewed as a matrix factorization task, where every element of the target matrix *y*^(*i,j*)^ is modeled as some form of a transformation of the corresponding hidden representations of the drug and cell line (Fig. 1a).

DEERS is a deep neural network-based recommender system. It consists of three major parts: drug autoencoder, cell line autoencoder and the subsequent feed-forward neural network. (Fig. 1b) (39, 40). The two autoencoder networks have the same architecture, with one 128-dimensional hidden layer in both encoder and the decoder with the rectified linear unit (ReLU) activation function, and the 10-dimensional hidden representation layer. The subsequent feed-forward network consists of a 20-dimensional input layer, followed by two hidden layers of length 512 and 256 with the ReLU activation. The regularization of the system is incorporated via the dropout with 0.5 probability, applied in the first, 512-dimensional hidden layer of the feed-forward network.

Consider a training data point consisting of original drug *i* and cell line *j* feature vector representations along with the corresponding response value, (**x**_*D*_^(*i*)^, **x**_*C*_^(*j*)^, *y*^(*i,j*)^). The input training data vectors are first passed into drug and cell line autoencoders, producing reduced, 10-dimensional vector representations (the hidden representations) (**z**_*D*_^(*i*)^, **z**_*C*_^(*j*)^) and reconstructed inputs 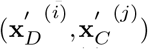 (Fig. 1b). The hidden representations **z**_*D*_^(*i*)^ and **z**_*C*_^(*j*)^ are then concatenated, forming a 20-dimensional vector, which serves as an input for the subsequent feed-forward neural network, which in turn computes the final response estimate *ŷ* ^(*i,j*)^ (Fig. 1b).

DEERS has three outputs and three main optimization goals: minimizing the differences between **x**_*D*_^(*i*)^ and 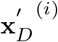 minimizing the differences between **x**_*C*_^(*j*)^ and 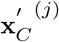, and minimizing the errors between *y*^(*i,j*)^ and *ŷ* ^(*i,j*)^. The incorporation of reconstruction errors causes the network to find informative representations of the input drug and cell line features. In addition, it is desired for the hidden dimensions to be independent. This enables the hidden representations to capture more information about the full input data and facilitates easier interpretations of the hidden dimensions. In the proposed model, it is achieved by minimizing the squared values in the off-diagonal entries of the drugs and cell lines covariance matrices in the latent space. All of the described optimization tasks are captured by a single cost function, which is iteratively minimized for each training batch to train the model:

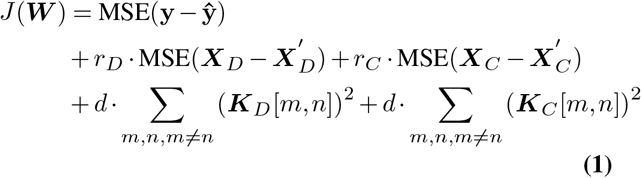

where *J* is the cost function, MSE denotes mean squared error, ***W*** is a set of the model parameters (weights), *r*_*D*_ is the real-valued weight of the drugs reconstruction error, ***X***_*D*_ is the drugs data matrix in the training batch, 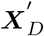 is the drugs data reconstruction matrix in the batch, *r*_*C*_ is a real-valued weight of the cell lines reconstruction error, ***X***_*C*_ is the cell lines data matrix in the batch, 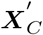 is the cell lines data reconstruction matrix in the batch, *d* is a weight of the dependence penalty, ***K***_*D*_ is the covariance matrix of drugs hidden representations in the batch, and ***K***_*C*_ is the covariance matrix of cell lines hidden representations in the batch, and ***K***[*m, n*] denotes the *m, n*-th entry of matrix ***K***.

Intuitively, the cost function weights *r*_*D*_, *r*_*C*_ and *d* control the contribution of the particular optimization task in the general optimization goal of the system. Setting all of these weights to zero would result in a network without decoding tasks and no dependence restrictions on the hidden dimensions of the drugs and cell lines.

### Compared models

We compare the proposed model to four other methods; two of which are based on traditional machine learning algorithms, while the other two are forms of matrix factorization.

In order to evaluate the traditional methods in a multi-task setting, where the data for all drugs and all cell lines are modeled at once, the traditional methods are used to predict drug response for the union of drugs and cell lines features. To this end, for every data point (**x**_*D*_^(*i*)^, **x**_*C*_^(*j*)^, *y*^(*i,j*)^), we first concatenate vectors **x**_**D**_^(*i*)^ and **x**_**C**_^(*j*)^, forming one 535-dimensional vector per drug-cell line pair. Applying this to all available data points produces a 52730 × 535 input data matrix ***X*** and the corresponding 52730-dimensional vector with true response values **y**. This data is used to train and evaluate two common machine learning algorithms: Elastic net (41) and XGBoost (42). The former is a linear model and the latter is a more complex, nonlinear model.

The compared matrix factorization models aim at solving a similar matrix-factorization type of problem (Fig. 1a) and can be seen as simpler or reduced versions of the proposed model. The first is a basic matrix factorization with side information method, reducing the dimension of the additional information about both drugs and cell lines using linear projections, and applying a dot product to produce the prediction of the response variable (here, the sensitivity of cell lines to drugs). We refer to this model as Lin MF (Fig. S1a). The basic architecture of this model is the same as the model applied by Yang *et al*. (26).

The second of the compared matrix factorization-based models is an non-linear extension of the basic model, where the dimensionality reduction is performed via one-layered autoencoders and data reconstruction is also taken into consideration (Fig. S1b). Similarly as in Lin MF, the final prediction is obtained by taking the dot product of the corresponding hidden representations, in contrast to the proposed DEERS model, where a separate feed-forward network is used to obtain the response estimate (Fig. 1b). We refer to this model as Autoen MF. To estimate the parameters of both Lin MF and Autoen MF we use gradient descent optimization implemented in Adam optimizer (43).

### Experimental setup and model training

In order to assess the performance of the considered models on the unseen cell lines, we construct the validation and test sets by first randomly selecting two sets of 100 unique cell lines each. We then extract the data points containing selected cell lines, producing the validation and test sets with ∼ 5000 drug-cell line pairs each. The rest of the pairs corresponding to the remaining 722 unique cell lines (with ∼ 42, 000 pairs) constitute the training set.

Before the training, the input cell line data were preprocessed by standard scaling of the continuous gene expression data so that every feature has zero mean and unit standard deviation, while binary coding variants and dummy encoded tissue types were unmodified. For the input drug data, all features were standardized in the same way as the gene expression. Since the GDSC AUC values are in the range of [0, 1], they were not scaled, while the log IC50 values were linearly preprocessed with min-max scaler to the [0, 1] range. Notably, all values necessary to perform each of the applied preprocessing schemes were calculated only on the training set and applied to the validation and test sets.

We use the training and validation sets in order to find the optimal set of hyperparameters, consisting of: network architecture, cost function weights *r*_*D*_, *r*_*C*_ and *d*, regularization type and learning rate. We establish the DEERS architecture as consisting of two-layer autoencoders, with 10-dimensional hidden representations (Fig. 1b). The subsequent feed-forward network has two hidden layers of size 512 and 256. The optimal cost function weights were set to *r*_*d*_ = 0.1, *r*_*C*_ = 0.25 and *d* = 0.1. As a regularization type, we use combination of dropout applied in the first hidden layer of the feed-forward network (Fig. 1b) and early stopping. With these hyperparameters fixed, for every split of the data (into the training, validation and test sets) we tune the learning rate, dropout rate and number of epochs for early stopping.

After all parameters are found, we use them to train the model using the union of training and validation sets, and apply the resulting model to the test set in order to assess the performance. We repeat this procedure five times with different cell lines in training, validation and test sets in order to improve the robustness of the results.

We adopt the similar methodology for the compared models, where we first tune the hyperparameters using training and validation sets, and then apply the final retrained model to the test set, using the same data splits for training, validation and testing for all models. In addition, we incorporate a simple data augmentation scheme, where we add a random gaussian noise with zero mean to the cell lines gene expression data and the corresponding AUC or IC50 values. The standard deviations of cell lines and response noise were 0.6 and 0.15, respectively. The augmentation was performed iteratively in every batch during training, tripling the original batch size. This data augmentation scheme was added for the two models involving autoencoders, i.e. both the Autoen MF and the DEERS model.

### Interpretation of hidden dimensions in DEERS

This analysis aims at an explanation of the model predictions from the biological standpoint. In order to incorporate all available data for model interpretation, we first re-train the model with all available 922 cell lines and 74 drugs, without excluding any cell lines, and using IC50 as a drug response metric.

The interpretation of the hidden dimensions concerns assigning a biological meaning to the individual dimensions of the hidden space. To this end, we first pass the input drugs and cell lines input representations into their corresponding, already trained autoencoders, producing a 10-dimensional representation for each 294-dimensional input data vector corresponding to a drug and a 10-dimensional representation for each 241-dimensional input data vector corresponding to a cell line, respectively.

### Associating input features with hidden dimensions

To compute the association of each input feature with each hidden dimension, we utilize the Integrated Gradients method (35), by computing the attributions between input features and the ten neurons constituting the hidden representation layers. This is performed separately for the drug and the cell line autoen-coders, and the attributions are averaged across the drugs and cell lines, respectively. As a result, we obtain drugs and cell lines feature-representation attribution matrices of size 294 × 10 and 241 × 10, respectively, where each entry is a score reflecting how much a given feature impacts the given variable in the hidden space. We then perform the row-wise hierarchical clustering on the resulting attribution matrices, grouping features associated with the same dimension together. The clustering was performed after normalizing the rows to unit norm, using the Ward linkage method and the Euclidean distance metric. This interpretability approach is applied separately for the 10 dimensions encoding the drugs and for the 10 dimensions encoding the cell lines.

### Associating biological processes with hidden dimensions encoding the cell lines data

In this interpretability analysis, we exploit the fact that the cell line autoencoder in DEERS is trained to reconstruct the data and to find low-dimensional representations that reflect the true properties of the analyzed cell lines. The produced hidden representations of the cell lines are organized into a 922 × 10 matrix ***Z***_*C*_, where every row *j* corresponds to a single cell line hidden representation, and every column *c* represents the values of a given hidden variable across all cell lines. Next, we examine the full genome-wide gene expression data of the full set 17,419 genes extracted from GDSC. In this way, this analysis goes beyond the restricted set of the modeled input 241 cell line features. Using this data, we construct a 922 × 17419 matrix ***G***_*C*_, where every row *j* corresponds to a single cell line gene expression profile, and every column *g* represents the expression values of a given gene across the examined cell lines. We then compute a 17, 419× 10 correlation matrix ***C***, where every entry ***C***[*g, c*] corresponds to Spearman correlation coefficient between *g*^*th*^ column of ***G***_*C*_ and *c*^*th*^ column of ***Z***_*C*_, i.e. the correlation between the expression of a given gene and a value of a given hidden dimension across 922 considered cell lines (Fig. 1c).

Given such correlation matrix ***C***, we create a ranked list of genes for every hidden dimension, where the ranking metric is the correlation coefficient of the genes with that dimension. The genes at the top and bottom of the ten resulting ranked lists are the ones that are most positively or negatively correlated with the corresponding dimensions, respectively. We then take the first and the last 1000 genes with corresponding correlation coefficients for every hidden dimension and run the GSEA Preranked analysis (44) against gene sets that are involved in specific biological processes as defined by the Biological Process GO Terms (Fig. 1c). The GSEA Pre-ranked is performed using the gseapy Python package (44– 46). We then extract the top 15 enriched terms with the smallest FDR value for every hidden dimension, which indicates the general biological mechanisms are most related to that dimension. Finally, we eliminate the redundant gene ontology terms using the Revigo tool (47), assigning the set of biological mechanisms to every dimension of the cell lines hidden space.

## Results

DEERS was developed with two aims in mind. One, to achieve state-of-the art predictive performance in predicting the response of cancer cell lines to kinase inhibitor drugs. Second, to identify the biological mechanisms that drive this response. Below, we evaluate the performance of DEERS in comparison to other models and conduct its interpretability analysis.

### Evaluation of the predictive performance of DEERS in comparison to other models

The predictive performance of DEERS is compared with four other methods. Two of those, Elastic net and XGBoost, are traditional, frequently used machine learning algorithms. Remaining two, referred to as Lin MF and Autoen MF, are versions of matrix factorization with side information (see Methods for a description of the compared models). In order to evaluate the performance of DEERS and other models on a test set containing responses of unseen cell lines, we first pass the drugs and cell lines input data to the model and obtain a table of predicted responses for each drug and cell line pair. Given such a table, we calculate the Pearson correlation and RMSE of the true to predicted responses across all drug-cell line pairs. In addition to such metrics calculated globally, we also group the previously described table, and calculate correlation (abbreviated corr.) and RMSE of true and predicted responses across pairs per given drug or cell line. To aggregate the per-drug and the per-cell line results, we take the median across the cell lines and drugs, respectively. The per cell line results mimic an envisioned clinical application of the model, where prediction of drug efficacy will be predicted for a new patient with specific tumor features, enabling a personalized medicine approach. This evaluation scheme yields six performance metrics per model (referred to as “Pairs RMSE”, “Pairs corr.”, “Per-drug RMSE”, “Per-drug corr.”, “Per-cl RMSE” and “Per-cl corr.”). These metrics are evaluated both for IC50 (Tab. 1) and AUC (Tab. 2). In general, IC50 as a prediction target was easier to learn than AUC. Indeed, in terms of correlation between predicted and true response values, better results are obtained by all models for IC50 than for AUC.

**Table 1.** Predictive performance of DEERS and compared models when using IC50 as a drug response metric. The presented values are averages of metrics taken across five experiments with different data splits, along with the corresponding standard deviations. The presented per-drug and per-cell line results are medians taken across all considered drugs and cell lines, respectively. The evaluated models are split into two categories: frequently used, traditional machine learning algorithms (T) and recommender system class (RS). Best results within a model category are highlighted with bold font, while the best results overall are underlined. Abbreviations: alg. - algorithm, corr. - correlation, cl - cell line, w/o inhib. profs. - without inhibition profiles.

|  | Alg. type. | Pairs RMSE | Pairs corr. | Per-drug RMSE | Per-drug corr. | Per-cl RMSE | Per-cl corr. |
| --- | --- | --- | --- | --- | --- | --- | --- |
| Elastic net | T | 0.09<br>±0.002 | 0.80<br>±0.007 | 0.08<br>±0.019 | 0.31<br>±0.155 | **0.08**<br><u>±0.002</u> | 0.84<br>±0.003 |
| XGBoost | T | **0.08**<br><u>±0.002</u> | **0.83**<br><u>±0.009</u> | **0.08**<br><u>±0.017</u> | **0.40**<br><u>±0.131</u> | **0.08**<br><u>±0.001</u> | **0.86**<br><u>±0.006</u> |
| Lin MF | RS | 0.09<br>±0.003 | 0.78<br>±0.011 | 0.09<br>±0.002 | 0.30<br>±0.040 | **0.08**<br><u>±0.002</u> | 0.85<br>±0.007 |
| Autoen MF | RS | 0.10<br>±0.003 | 0.76<br>±0.012 | 0.09<br>±0.004 | 0.24<br>±0.039 | 0.09<br>±0.004 | 0.84<br>±0.004 |
| DEERS w/o inhib. profs. | RS | 0.09<br>±0.002 | 0.80<br>±0.012 | **0.08**<br><u>±0.002</u> | **0.38**<br><u>±0.047</u> | **0.08**<br><u>±0.002</u> | 0.84<br>±0.003 |
| DEERS | RS | **0.08**<br><u>±0.001</u> | **0.82**<br><u>±0.004</u> | **0.08**<br><u>±0.002</u> | 0.37<br>±0.019 | **0.08**<br><u>±0.001</u> | **0.86**<br><u>±0.006</u> |

**Table 2.**
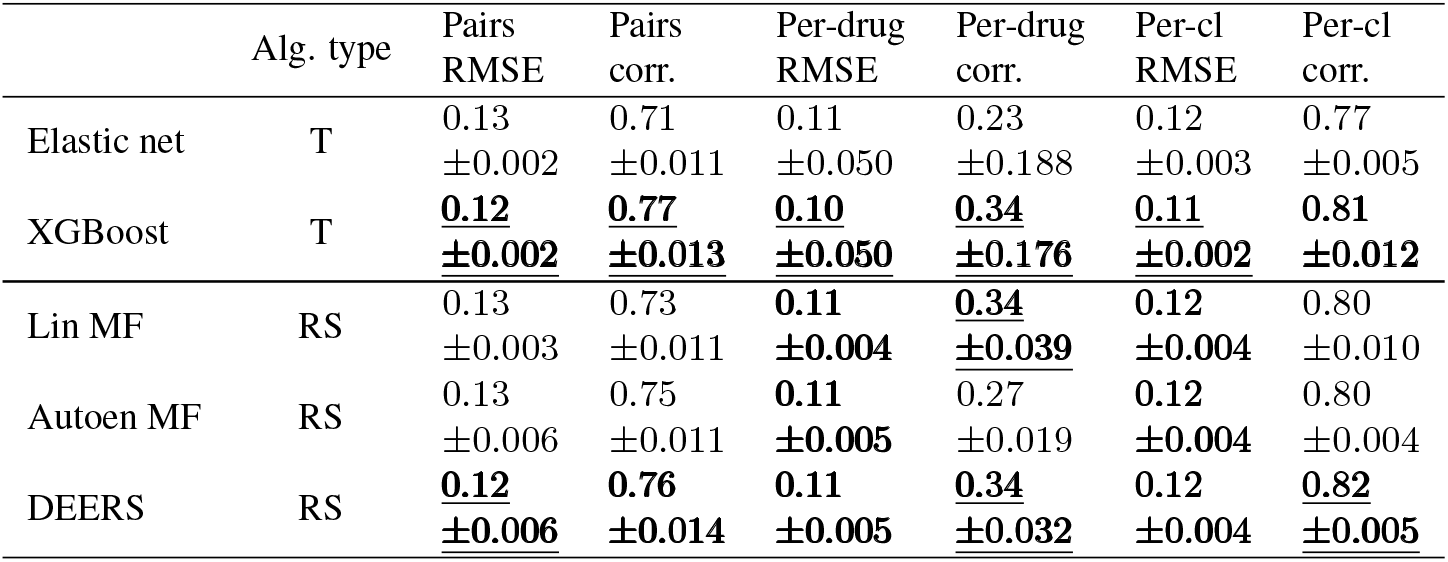
Predictive performance of DEERS and compared models when using AUC as a drug response metric. Table columns and formatting the same as in Tab. 1.

|  | Alg. type | Pairs RMSE | Pairs corr. | Per-drug RMSE | Per-drug corr. | Per-cl RMSE | Per-cl corr. |
| --- | --- | --- | --- | --- | --- | --- | --- |
| Elastic net | T | 0.13<br>±0.002 | 0.71<br>±0.011 | 0.11<br>±0.050 | 0.23<br>±0.188 | 0.12<br>±0.003 | 0.77<br>±0.005 |
| XGBoost | T | **0.12**<br><u>±0.002</u> | **0.77**<br><u>±0.013</u> | **0.10**<br><u>±0.050</u> | **0.34**<br><u>±0.176</u> | **0.11**<br><u>±0.002</u> | **0.81**<br><u>±0.012</u> |
| Lin MF | RS | 0.13<br>±0.003 | 0.73<br>±0.011 | **0.11**<br><u>±0.004</u> | **0.34**<br><u>±0.039</u> | **0.12**<br><u>±0.004</u> | 0.80<br>±0.010 |
| Autoen MF | RS | 0.13<br>±0.006 | 0.75<br>±0.011 | **0.11**<br><u>±0.005</u> | 0.27<br>±0.019 | **0.12**<br><u>±0.004</u> | 0.80<br>±0.004 |
| DEERS | RS | **0.12**<br><u>±0.006</u> | **0.76**<br><u>±0.014</u> | **0.11**<br><u>±0.005</u> | **0.34**<br><u>±0.032</u> | **0.12**<br><u>±0.004</u> | **0.82**<br><u>±0.005</u> |

With IC50 as the response variable, the DEERS model mostly outperforms or at least performs similarly well as the other two matrix factorization-based models with regard to all of the six performance metrics (indicated by bolded values in Tab. 1). For IC50, the XGBoost outperforms the other traditional method, Elastic net, in all performance measures. This indicates that nonlinear models are needed to capture the dependence of IC50 on drug and cell line features. DEERS and XGBoost achieve comparable evaluation results (with the best model according to each evaluation metric marked in red in Tab. 1). In particular, DEERS obtains a high Pearson correlation coefficient r=0.82, calculated on all drug-cell line pairs in the test set. Moreover, the median per cell line correlation of r=0.86 indicates that DEERS achieves the state-of-the-art performance in predicting cell line responses to drugs, which most closely resembles the hypothetical clinical setup. Notably, compared to per-cell line correlation, all models obtain relatively poor results in terms of per-drug correlation. This may be due to the fact that our input data is asymmetric as it covers much fewer drugs (74) than cell lines (922). In the case of AUC as the response variable, the comparison of model performance yields similar results as the IC50. Here again DEERS outperforms the other two matrix factorization-based methods, while from the two traditional methods XGBoost performs better than Elastic net (Tab. 2). Overall, the performance of DEERS is very similar to XG-Boost. For AUC, the DEERS achieves r=0.76 Pearson correlation coefficient calculated on all drug-cell line pairs in the test set. For the per-cell line results, the median correlation across the unseen cell lines is r=0.82, constituting the best result by a small margin.

### Evaluation of the added value of inhibition profiles

In order to quantify the benefit of incorporating inhibition profiles of the drugs, we estimated the performance of DEERS with drug putative targets as drug input data. This model is referred to as “DEERS without inhibition profiles” in Tab. 1. To this end, the reduced drug features were defined by a binary matrix with 74 rows corresponding to kinase inhibitors and 92 drug targets, and entries 1 if the drug has the gene as target and 0 otherwise. With this alternative drug input data and IC50 as a target variable, we evaluated DEERS using the same procedure as previously, with all hyperparameters besides learning and dropout rates unchanged. Learning and dropouts rates were tuned using validation set in the same manner as before. DEERS with inhibition profiles out-performs DEERS with binary targets in 3 evaluation metrics (Pairs RMSE, Pairs corr., Per-cl corr.), achieves the same results in 2 metrics (Per drug RMSE and Per-cl RMSE) and slightly underperforms in Per-drug corr. metric. The improvement in Pairs RMSE, Pairs corr., Per-cl corr. metrics constitutes 11.1%, 2.5%, and 2.4% relative increase, respectively.

### Evaluation on an independent dataset

In order to estimate the performance of DEERS on other data than cell line sensitivities from GDSC, we extracted drug sensitivity data from the Cancer Cell Line Encyclopedia (CCLE) (2) project. Next, we constructed a dataset consisting of an intersection between the our analyzed dataset (containing data for 74 drugs derived from GDSC for kinase inhibitors), and the CCLE dataset in terms of cell lines and drugs, along with corresponding, min-max-scaled CCLE IC50 values. The data regarding the intersection between GDSC and CCLE, as well as CCLE IC50 values were extracted using the PharmacoDB package (48, 49). The resulting dataset contained 351 common cell lines and 5 common drugs (Crizotinib, Lapatinib, PD0325901, PLX-4720 and Sorafenib), constituting 1747 pairs in total. The cell lines and drugs were described by the same features as in the original GDSC dataset. We next used the GDSC data corresponding to the remaining 571 cell lines that are not present in the CCLE-GDSC intersection dataset and all 74 drugs to train DEERS. From those 571 cell lines of GDSC, 50 were randomly chosen to construct the validation dataset for tuning the learning and dropout rates (see Methods). We then re-trained the model with the best hyperparameters on all 571 cell lines and applied it to the CCLE-GDSC intersection dataset, obtaining IC50 predictions for unseen cell lines. It is important to note that the maximum obtainable correlation between the model predictions and the true IC50 values in the intersection dataset in this experiment is 0.53, defined by the correlation between the true IC50 values in the CCLE dataset and the true IC50 values in the GDSC for these cell line-drug pairs. Given this upper bound, the obtained correlation result of 0.40 is relatively high.

Taken together, these results demonstrate that thanks to its deep neural network-based recommender system architecture and utilization of informative drug features, DEERS obtains state-of-the art performance in predicting cell lines sensitivity to drugs in a multitask setup. In contrast to the other well performing model, XGBoost, however, DEERS obtains highly informative reduced-dimension representations of the cell line and drug features, respectively. This aspect of the model is discussed below.

### Attributions between input features and hidden dimensions using neural network analysis

As the first step of the DEERS model interpretability analysis, we computed the attributions between the input features and the hidden dimensions using Integrated Gradients (see Methods). Next, we performed hierarchical clustering of the resulting attribution matrix, in which the rows were the features, and columns were the hidden dimensions. The clustering identifies well-defined groups of features associated with each specific hidden dimension (Fig. 2). There is very little overlap between feature groups for both drugs and cell lines, indicating that hidden dimensions are independent in terms of which features affect them the most. This independence effect is also evident when we compare the covariance matrices of drugs and cell lines represented in a hidden space, when dependence penalty was incorporated and not incorporated into the overall cost function (Fig. S2). The number of drug input features associated with a single hidden dimension ranges from 20 for dimensions 2 and 8, to 44 for dimension 1. For the cell lines, this number ranges from 11 for hidden dimension 7 and 44 for hidden dimension 1.

**Fig 2.**
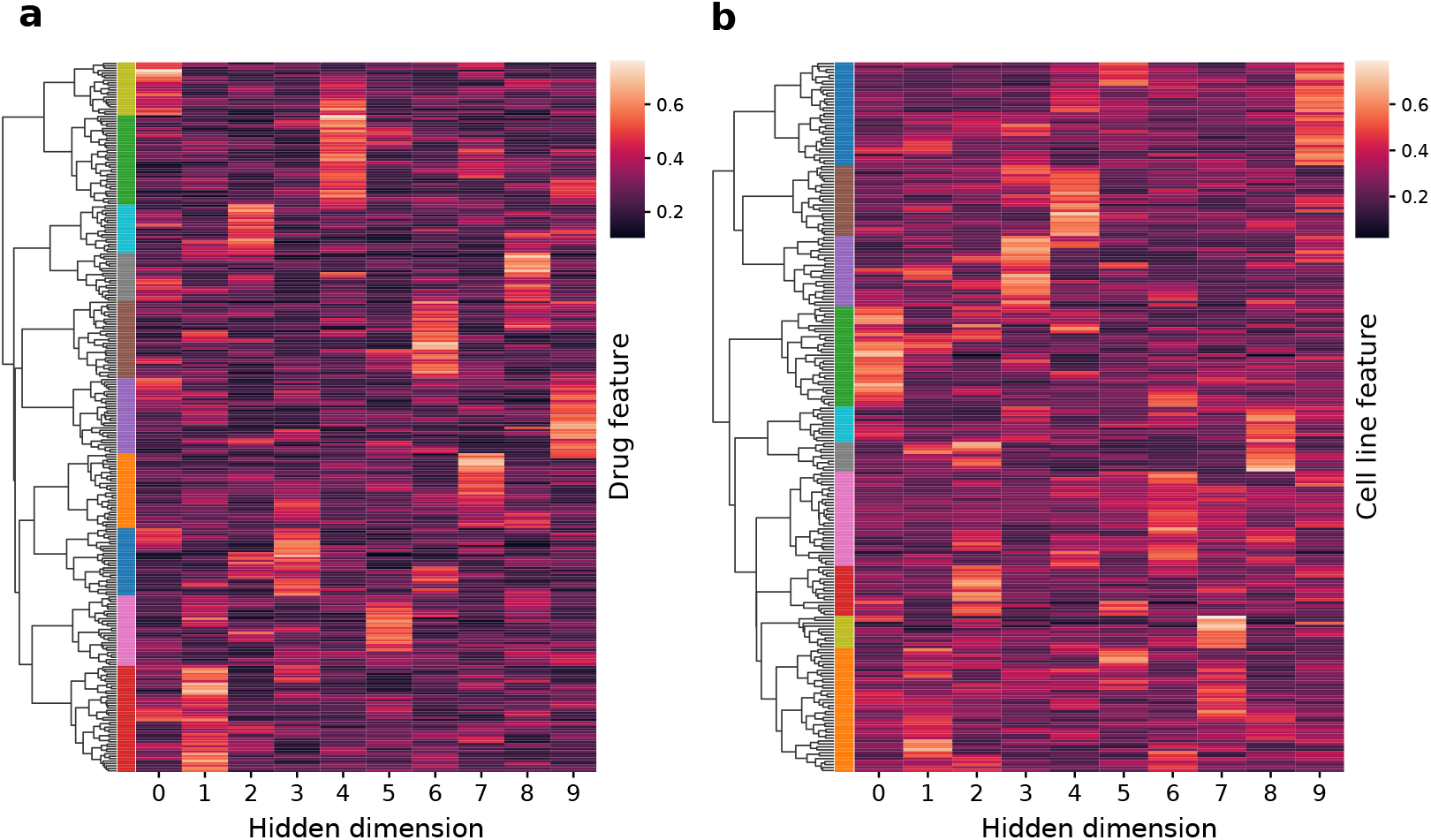
Clustermaps of attribution scores between input features and hidden dimensions for (a) drug autoencoder and (b) cell line autoencoder. Color reflects the importance of a given feature for a given hidden dimension. The vertical color bars next to the dendrograms represent the cluster assignment of groups of features.

The exact groups of features corresponding to hidden dimensions of the drug autoencoder and to hidden dimensions of the cell line autoencoder are listed in tables S1 and S2, respectively. As an example, in the drugs case, hidden dimension 3 is associated with the inhibition of a group of kinases (BRSK1, CAMK2B, CDK3, CDK5, CDK16, CHEK1, DCLK1, DRAK1, ERBB4, ERK5, FRK, HCK, LIMK2, MAPKAPK2, MK14, MLK1, MP2K6, NDR2, PCTK3, PIM1, PIM2, PLK1, PRKR, TYRO3, VGFR1, VGFR2, YANK3 and YSK1). For the cell lines example, hidden dimension 3 is associated with the expression of genes *BLK, BRSK1, BTK, CSK, DDR1, EGFR, EPHA2, FGFR2, GAK, LCK, MET, NLK, NUAK1, NUAK2, PIM2, PLK2, RIOK1* and *ZAP70*, as well as one-hot encoded tissue indication features corresponding to leukemia, lung NSCLC, lymphoma, myeloma, pancreas and urogenital system.

### Linking hidden dimensions to the general biological mechanisms

In the next step of the interpretability analysis, we associate each hidden dimension of the cell line autoencoder with a biological process. To this end, for each hidden dimension and each gene, we correlate the values of the hidden dimension with the expression values of the gene across cell lines. For a given hidden dimension, the obtained correlations are then ranked and we apply gene set enrichment analysis (GSEA) to identify biological processes positively or negatively correlated with that dimension (Fig. 1c). Importantly, this analysis links the dimensions to all genes measured in the cell lines, that is, also to the genes outside of the cell line features used in the model (see Methods for a full description of this analysis). Here, we run the GSEA considering the gene-sets included in the Gene Ontology Biological Processes. The analysis and subsequent filtering of redundant terms yield a final set of GO terms for each dimension of the hidden space of the cell line autoencoder (Fig 3). We identify 67 GO terms in total, many of which are related to cancer (e.g. DNA replication, regulation of cell cycle process, regulation of angiogenesis). The number of enriched terms per dimension varies from 6 to 13. The majority of enrichment scores (67%) are positive, which indicates that they are positively correlated with that dimension. Conversely, the negatively signed FDR value implies that the given term is negatively correlated. Markedly, the sets of enriched terms almost do not overlap between the dimensions, indicating the independence of the dimensions in terms of their associated biological mechanisms. Out of 67 terms, only 12 are associated with more than one hidden dimension, from which 10 are associated with two dimensions.

**Fig 3.**
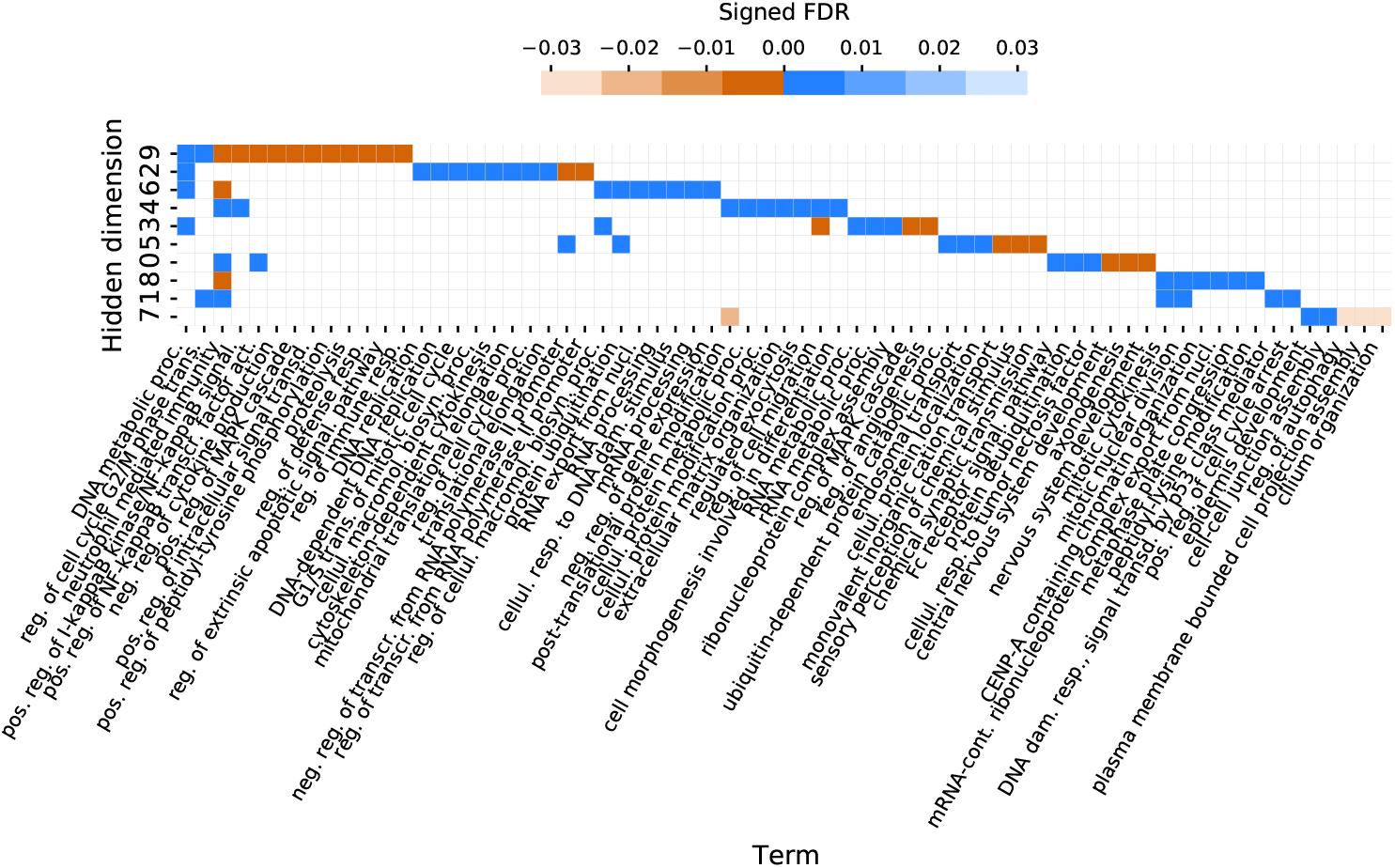
Heatmap reflecting Biological Process GO terms enriched in hidden dimensions of the cell line autoencoder. Negatively and positively signed FDR values correspond to negative and positive enrichment scores, i.e. negative and positive side of the ranked list, respectively. Hidden dimensions are sorted by the number of enriched terms. Abbreviations: proc. – process, reg. – regulation, resp. – response, neg. – negative, pos. – positive, transcr. – transcription, transd. – transduction, mRNA-cont – mRNA-containing, dam. – damage, nucl. – nucleus, macromol. – macromolecule, biosyn. – biosynthetic, cellul. – cellular, signal. – signaling, act. – activity, trans. – transition.

When inspecting the heatmap (Fig. 3), we identify groups of biological mechanisms associated with specific hidden dimensions. For example, hidden dimension 2 is mainly linked with DNA replication and cell cycle, as terms enriched in it include: DNA replication, DNA-dependent DNA replication, G1/S transition of mitotic cell cycle and regulation of cell cycle process. Dimension 4 is related to protein metabolism (post-translational protein modification, cellular protein metabolic process, cellular protein modification process), while dimension 3 is connected with DNA and RNA metabolism (DNA metabolic process, RNA metabolic process, rRNA metabolic process) and known cancer-related processes like regulation of MAPK cascade and regulation of angiogenesis. Other such terms include regulation of extrinsic apoptotic signaling pathway (dimension 9), cellular response to DNA damage stimulus (dimension 6), cellular response to tumor necrosis factor (dimension 0) and DNA damage response, signal transduction by p53 class mediator (dimension 1). Interestingly, some of the terms are not commonly linked to cell cycle or other processes related to oncogenesis, e.g. for dimension 8 the set of enriched terms includes central nervous system development, nervous system development and axonogenesis. This analysis provides a form of interpretation of hidden dimensions from the biological standpoint and facilitates a better understanding of the model prediction based on cell lines hidden representations. Overall, the obtained list of biological processes reflects the repertoire of common biological mechanisms that are affected by the analyzed kinase inhibitors in the set of analyzed cell lines, and as a general summary can only be obtained from such a multitask learning model as DEERS.

### Case studies

We further focus the analysis on three case studies, showing how the model predictions and true responses can be explained and interpreted for individual drugs and features. For this purpose, we examine three specific compounds: the pan-CDK inhibitor PHA-793887, the ALK/CDK7 inhibitor XMD14-99 and the BRAF inhibitor Dabrafenib (Fig. 4). First, we establish which features are most important for the model prediction given the input data for a particular compound. To this end, we calculate the attributions between input features and the final output layer of the model using the Integrated Gradients method (35). The attributions are first computed separately for each cell line and IC50 as the response variable, and next summarized by averaging over all cell lines. Second, for each compound we display the cell lines in two chosen dimensions of the hidden space of the cell line autoencoder, and color them by their IC50 response to the compound. In this way, we identify such regions in this space that are correlated with sensitivity to the compound. Finally, we explore in detail how well one chosen hidden dimension correlates with the true response and we list the biological processes that are associated with that dimension (as per analysis in Fig. 3). Altogether, the case studies identify such features and hidden dimensions that are important for modeling the response, and such biological processes that are important for the action of the three analyzed drugs.

**Fig 4.**
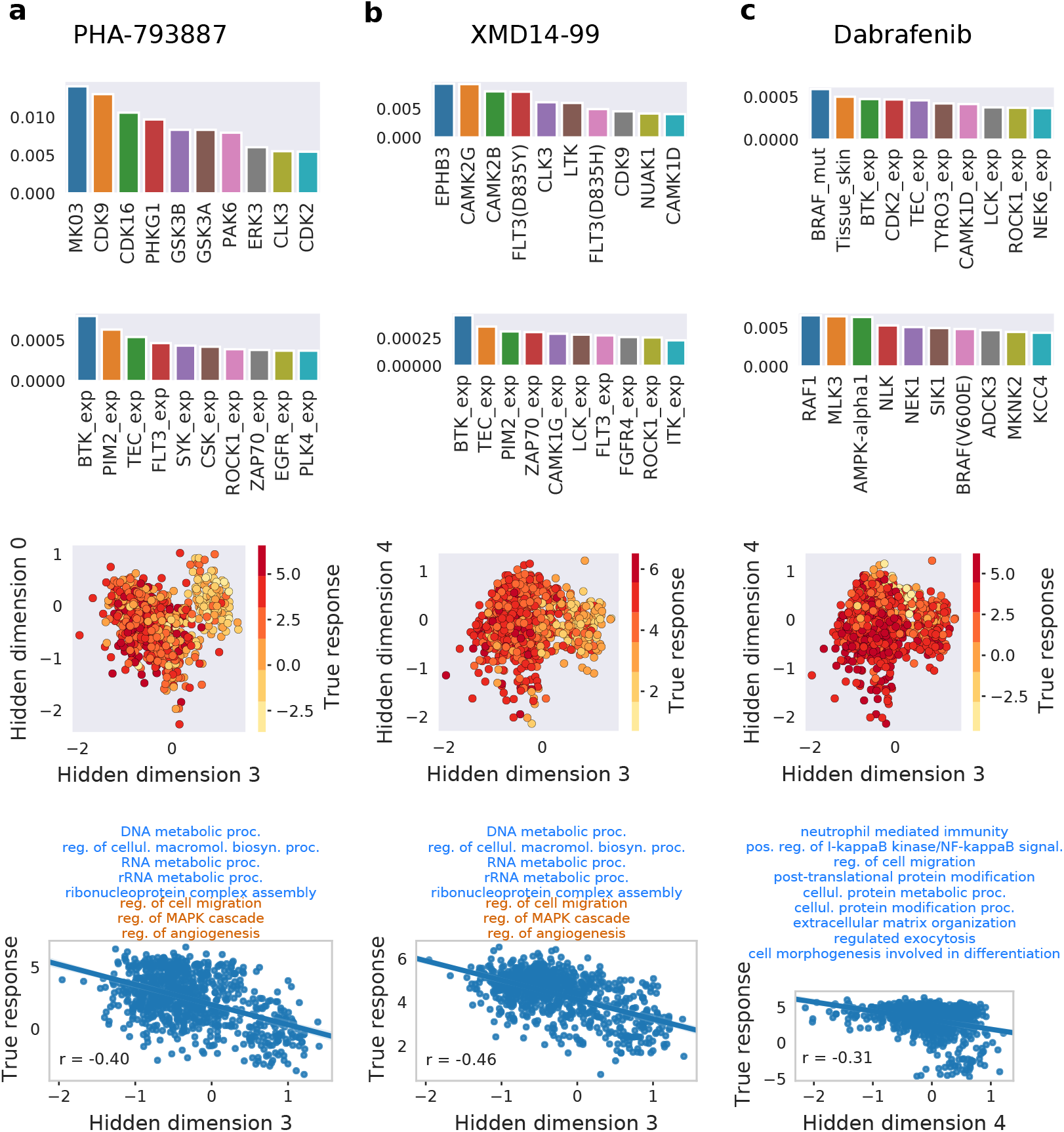
Case studies corresponding to compounds: (a) PHA-793887, (b) XMD14-99 and (c) Dabrafenib. Top row panels: top ten most important drug features for a response prediction by the model, derived using Integrated Gradients. Second row panels: top ten most important cell lines features for a model’s response prediction. Feature names abbreviations: exp. – expression, mut. – mutation. Third row panels: cell lines plotted using two chosen hidden dimensions of the cell line hidden space, colored by the true log IC50 values (shown for those cell lines that were screened against the presented drug). Bottom row panels: scatter plots of true log IC50 values w.r.t. hidden dimension most correlated with the response for a given drug. The presented r values are the Spearman correlation coefficients. Text on top shows which GO terms are enriched in a considered hidden dimension, following Fig. 3, where blue and brown colors correspond to positive and negative enrichment, respectively. See Fig. 3 for term names abbreviations.

PHA-793887 is an inhibitor of multiple cyclin dependent kinases (CDK) with activity against CDK2, CDK1 and CDK4 (50). According to the attribution analysis, the activity against CDK is reflected in the most informative drug features, where CDK2 kinase belongs to the most important drug features for prediction (Fig. 4a, top row panel). Interestingly, cell line features corresponding to CDK family do not obtain top attribution values. Instead, the highest average attributions are associated with the expression of *BTK, PIM2* and *TEC* genes, suggesting that their activity in the cell lines is important for PHA-793887 action (Fig. 4a, second row panel). Representing cell lines in two dimensions (by hidden dimensions 3 and 0) identifies a region corresponding to a good response of PHA-793887 (Fig. 4a, third row panel). This validates that that in general the hidden dimensions well represent the cell line data and that in particular these two hidden dimensions well capture the cell line response to PHA-793887. However, most of the cell lines response variance can be explained using hidden dimension 3 alone, which is negatively correlated with the true response (Pearson correlation r = −0.40; Fig. 4a, bottom row panel). The biological process terms enriched for different dimensions, visualized in Fig. 3, can provide the meaning behind these dimensions. Analysing the processes associated with the most informative dimension 3 can shed the light on the way the response to PHA-793887 is conveyed in the cell lines. The hidden dimension 3 is associated with eight biological processes, five of them positively (DNA metabolic process, regulation of cellular macromolecule biosynthetic process, RNA metabolic process, rRNA metabolic process, ribonucleoprotein complex assembly) and three of them negatively (regulation of cell migration, regulation of MAPK cascade, regulation of angiogenesis). Since the GSEA analysis is performed using gene expression data, large values of hidden variable 3 (which correspond to a better response) implicitly indicate the over-expression of genes associated with the five positively enriched terms, whereas the over-expression of genes related to three negatively enriched terms can indicate poor response.

According to GDSC annotations, XMD14-19 targets include ALK, CDK7, LTK and others. The known target LTK is listed as one of the most important drug features (i.e., with a large attribution; Fig. 4b, top row panel). Cell line features with top attributions for XMD14-19 strongly overlap with those related to PHA-793887 (Fig. 4b, second row panel). In particular, the top three to features are exactly the same (expression of *BTK, PIM2* and *TEC* genes), indicating some similarity between these two drugs. Hidden dimensions 3 and 4 allow to visualize regions in the cell line hidden space with distinctive responses (Fig. 4b, third row panel). Similarly to PHA-793887, hidden dimension 3 carries the most information about cell lines response to XMD14-99 and thus we can conclude that the same biological processes may be associated with the response to these two drugs (Fig. 4b, bottom row panel).

In the case of Dabrafenib, both drug and cell line feature sensitivities are consistent with its design and clinical usage. Dabrafenib is a selective inhibitor of mutant BRAF kinase, approved by the FDA for the treatment of metastatic melanoma with mutant BRAF(V600) (51, 52). Accordingly, the two most important cell line features are *BRAF* mutation and skin tissue indicator (Fig. 4c, second row panel). The inhibition of BRAF also emerges among the most informative drug features, though being preceded by the inhibition of RAF1, MLK and AMPK (Fig. 4c, top row panel). The hidden dimensions 3 and 4 capture significant information about cell lines response (Fig. 4c, third row panel), with the hidden dimension 4 being the sole good indicator of the Dabrafenib efficacy (Fig. 4c, fourth panel). The hidden dimension 4 has nine positively enriched biological process terms associated with it (Figures 3; 4c, bottom row panel). Thus, we conclude that a better response to Dabrafenib corresponds to the over-expression of genes involved in: neutrophil mediated immunity, positive regulation of I-kappaB kinase/NF-kappaB signaling, regulation of cell migration, post-translational protein modification, cellular protein metabolic process, cellular protein modification process, extracellular matrix organization, regulated exocytosis, and cell morphogenesis involved in differentiation.

### Associating biological processes to all of the analyzed drugs

In the final step of the interpretability analysis, we associate biological processes to all of the analyzed drugs. This analysis is based on the idea behind the bottom panels of Fig. 4. Just like for PHA-793887, XMD14-99 and Dabrafenib, we can calculate the correlation coefficient between the response profile for a given drug and a given hidden dimension across cell lines, for each of 74 drugs and each of 10 hidden dimensions (Fig. 1d). This calculation yields a 74 × 10 matrix, in which each entry represents a Spearman correlation coefficient for a given compound and hidden dimension. We then utilize the associations between hidden dimensions and biological processes presented in Fig. 3 in order to connect drugs to biological processes. For a given drug and process, we first establish which hidden dimension is enriched for that process. Next, we assign a corresponding correlation coefficient between the drug response and the dimension to the drug and process pair. If more than one hidden dimension is enriched for the process, we take the average of the corresponding correlations. This analysis produces a 74× 67 drug-process matrix, where each entry is a correlation coefficient indicating how important a given biological process is for driving the response of a cell line to a given drug. We divide this matrix into five sub-matrices by the main target pathways of the drugs: RTK signaling, PI3K/MTOR signaling, ERK MAPK signaling, Cell cycle, and Others. Finally, we perform the row-wise hierarchical clustering of each such drug-process sub-matrix in order to group drugs by the similarity of processes that drive their efficacy (Fig. 5). The obtained clustermaps clearly indicate such processes that are shared among drugs targeting the same pathways, as well as point at their differences, some of which are related to the particular gene targets.

**Fig 5.**
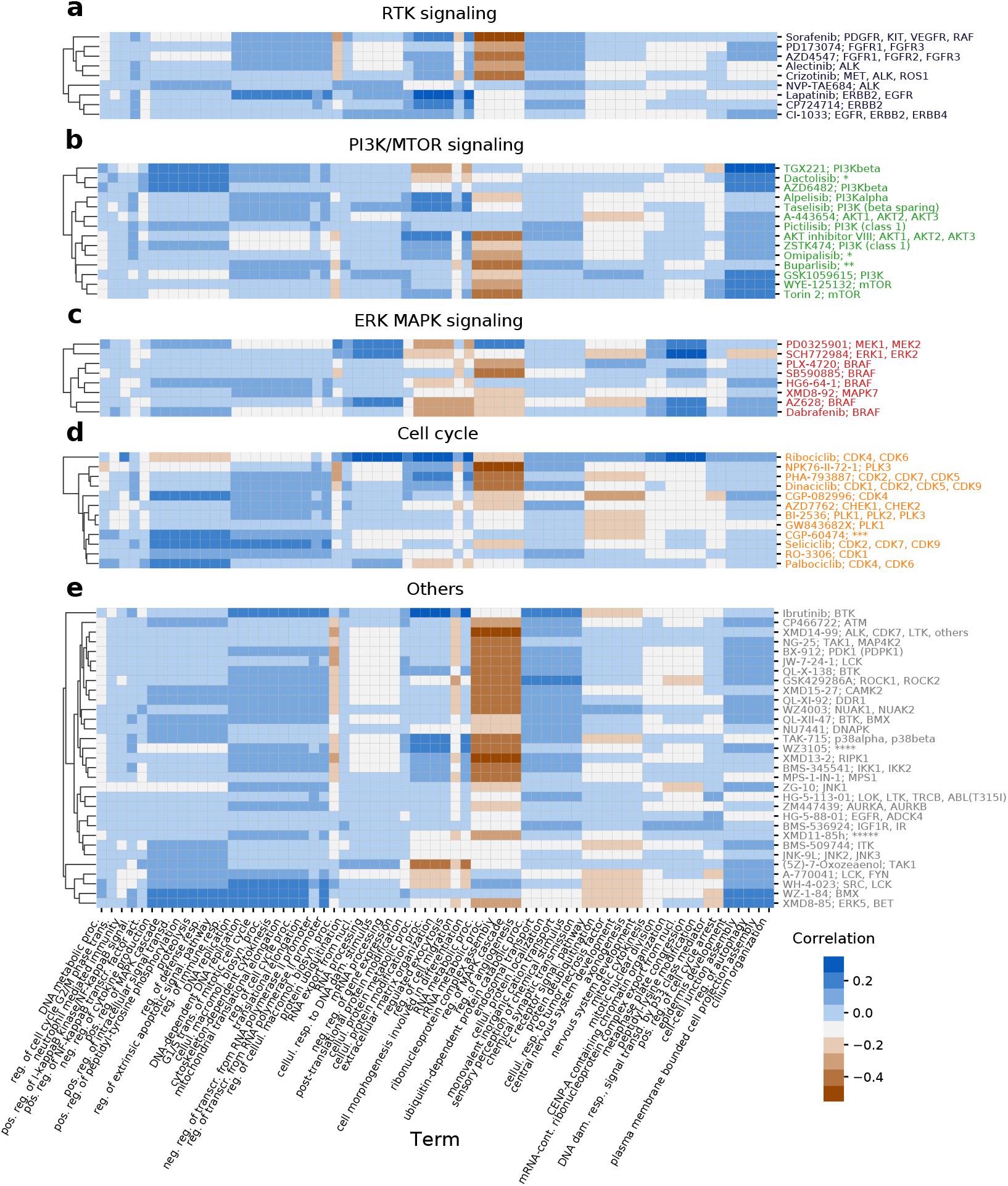
Associations between cell lines biological processes (horizontal axes) and all of the 74 analyzed drugs (vertical axes), plotted separately for (a) RTK signaling, (b) PI3K/MTOR signaling, (c) ERK MAPK signaling, (d) Cell cycle and (d) Others target pathways. First, for every drug, the Spearman correlation coefficient between every cell line hidden dimension and the response is computed, as shown in Fig. 4. These correlations are then assigned to biological processes associated with a given hidden dimension (see Fig. 3). If more than one hidden dimension is related to a process, an average of correlation is taken and assigned to the process. Drugs were hierarchically clustered using the Euclidean distance and average linkage within a given target pathway category. The horizontal axis is shared for all panels. The vertical axis tick label is formatted as: Drug Name; Putative Targets. Drugs target pathways and putative targets are taken from GDSC annotations. The labels are color-coded by the target pathway. For some drugs, putative targets have not been listed for readability. Those targets are: * – PI3K (class 1), MTORC1, MTORC2, ** – PI3Kalpha, PI3Kdelta, PI3Kbeta, PI3Kgamma, *** – CDK1,CDK2,CDK5,CDK7,CDK9, PKC, **** – RC, ROCK2, NTRK2, FLT3, IRAK1, others, ***** – BRSK2, FLT4, MARK4, PRKCD, RET, SRPK1. The color scale of correlation heatmaps is the same for all categories. See Fig. 3 for term names abbreviations.

All drugs targeting the RTK signaling pathway (Fig. 5a) are positively correlated with a large group of processes related to DNA replication and cell cycle (DNA replication, DNA-dependent DNA replication, G1/S transition of mitotic cell cycle, cellular macromolecule biosynthetic process, cytoskeleton-dependent cytokinesis, mitochondrial translational elongation, regulation of cell cycle process, translational elongation, negative regulation of transcription from RNA polymerase II promoter, regulation of transcription from RNA polymerase II promoter) as well as processes related to transport and sensory perception (ubiquitin-dependent protein catabolic process, processes endosomal transport, cellular protein localization, monovalent inorganic cation transport, sensory perception of chemical stimulus and chemical synaptic transmission). These drugs, however, visibly divide into two distinct groups with respect to a group of processes related to RNA metabolism and regulation of MAPK cascade and angiogenesis (RNA metabolic process, rRNA metabolic process, ribonucleoprotein complex assembly, regulation of MAPK cascade, regulation of angiogenesis). This difference is the reflection of putative targets of the drugs; drugs which have ERBB2 or EGFR as the putative targets are not correlated with these processes, while remaining drugs are negatively correlated with them. In general, the RTK signaling drugs with shared target genes have similar associated processes and cluster together.

Drugs targeting the PI3K/MTOR signaling pathways generally share very similar profiles of association with biological processes (Fig. 5b). All of the drugs in this category have consistent positive correlation with five processes (epidermis development, cell-cell junction assembly, regulation of autophagy, plasma membrane bounded cell projection assembly and cilium organization), which distinguishes PI3K/MTOR signaling from other drug categories. There are, however still some processes which tend to be correlated only with a subset of drugs. For example, seven drugs (AKT inhibitor VIII, ZSTK474, Omipalisib, Buparlisib, GSK1059615, WYE-125132 and Torin 2) exhibit stronger negative correlation with five previously listed processes related to RNA metabolism and regulation of MAPK cascade and angiogenesis, while others do not. The associations with the foregoing processes seem to be more prevalent in drugs which have the mammalian target of rapamycin (mTOR) kinase among their putative targets in addition to phosphoinositide 3-kinases (PI3Ks), with the exception of Dactolisib. Notably, two drugs presumably targeting solely mTOR (WYE-125132, Torin 2) have very similar association profiles across all 67 processes. Another considerable group of processes with relatively high correlation in the PI3K/MTOR signaling category consists of eight processes: negative regulation of cytokine production, positive regulation of MAPK cascade, positive regulation of intracellular signal transduction, positive regulation of peptidyl-tyrosine phosphorylation, proteolysis, regulation of defense response, regulation of extrinsic apoptotic signaling pathway and regulation of immune response. Two out of three drugs associated with these processes (TGX221 and AZD6482) target solely PI3Kbeta. Associations with the listed eight processes are also noticeable in the cell cycle category (Fig. 5d), specifically for CGP-082996, CGP-60474, Seliciclib and Palbociclib. In general, these results can serve as a validation and explanation of the model predictions, as well as provide insights regarding the drugs mechanisms of action and drivers of the cell lines response. Considering drug-biological process associations within a certain drug category enables insights into drugs action on a more general level than putative targets or target pathway information alone.

## Discussion and conclusions

In this work, we propose a deep neural network recommender system-based approach to the problem of kinase inhibitor sensitivity prediction based on side information about drugs and cancer cell lines. The proposed model, DEERS, combines dimensionality reduction of the cell line and drug features using autoencoders and neural network-based prediction based on the obtained hidden representations. The modeled drug features are the strengths of inhibition of kinases by the drugs. The cell line features include expression and mutation calls for the same kinases in cancer cell lines, complemented by primary tissue type of origin for the cell lines. To our knowledge, this type of modeling using these types of input data has not been applied before to predict sensitivity to kinase inhibitors.

Our focus on modeling kinase inhibitors is motivated by the fact that binding profiles across kinases represent exquisite data to characterize such drugs. Alternative information about drugs could be the list of specific known drug targets. In contrast to continuous and rich data about kinase inhibition, however, annotations of known targets are relatively incomplete. The quality of the kinase binding data that we used is assured by a standardized assay platform interrogating a large number of kinases. Therefore, off-target inhibition effects are most likely captured completely. An alternative could be to use information on which signaling pathways are affected by a drug since this information is often provided in drug databases. However, clearly the information about target pathways is only high-level, less detailed than using kinase binding data, and suffers from incomplete understanding about the complete set of pathways that a drug effects in different cellular contexts.

The DEERs model aims at two goals: 1) high predictive performance and 2) outstanding model interpretability. Our analysis constitutes a thorough comparative assessment of model performance, evaluating both traditional and variants of matrix factorization-based methods. Out of the two traditional models, XGBoost achieves better results than Elastic net, indicating that accounting for non-linear interactions among features is crucial for prediction performance. DEERS outperforms the other two matrix factorization-based approaches, Lin MF (basic matrix factorization model) and Autoen MF (a model using autoencoders for dimensionality reduction and a dot-product for combining the reduced data to make prediction). We observe little difference in performance between the Lin. MF model and Autoen. MF (Tables 2, 1), although, importantly, the latter has a more difficult optimization goal. Indeed, similar to DEERS, Autoen MF reconstructs the input features from the reduced representations. The advantage of DEERS over these two MF-based models is most likely caused by the incorporation of the feed-forward network which combines the hidden representations, instead of the simple dot product. Compared to the dot product, which only considers element-wise product, these additional feed-forward network layers allow the system to adjust the weights after the data encoding step and to estimate a more complex function that maps from the hidden representations to the response. Importantly, despite the more complex mapping, the hidden dimensions in the DEERS model are still clearly indicative of the true response in some cases. Across all compared models, both DEERS and XGBoost show top and very similar performance. In contrast to XG-Boost, however, DEERS is easier to interpret, as it provides highly informative 10-dimensional representations of the input cell lines molecular setup and the drug features.

Extensive interpretability analysis demonstrates that the 10 hidden dimensions of the drug and cell line autoencoders seem to capture the majority of important information for both drugs and cell lines. The results imply mutual independence of hidden dimensions (Fig. 2, 3, S2) and also suggest that the hidden representations are representative of the drug and cell line input data. In particular, hidden dimensions 3 and 4 show as most relevant for driving the cell lines drug response for the majority of drugs (Fig. 3, 5), as demonstrated by the presented examples (Fig. 4, bottom panels). The correlation analysis of genome-wide cell lines features and hidden representations, combined with GSEA, helps to provide biological meaning to the hidden dimensions (Fig. 3). The same analysis performed using the restricted set of kinase- and tissue type-related 241 cell lines features that are used to train the models would have resulted in the bias towards GO terms or pathways related to protein kinases in general. Instead, using genome-wide gene expression helps to identify the enriched terms which are not influenced by the choice of features in the training data and spanning a broader range of biological processes. Moreover, this methodology is potentially very versatile, as different drugs and cell lines properties outside of the training data can be correlated, and different gene set libraries can be queried for enrichments. We show that combining the influence of distinct hidden dimensions for drug response, and biological processes associated with the hidden dimensions constitutes a framework which can directly explain drug response by concrete biological mechanisms. Such a map facilitates the easier explanation of drugs mechanisms of action and can potentially identify the new, unexpected ones (Fig. 5). Overall, this study shows how data encoding combined with the series of analyses can help to increase the interpretability even in the case of deep neural network recommender models, while maintaining the complex nature of such systems.

