## Supplementary Information for "Interpretable deep recommender system model for prediction of kinase inhibitor efficacy across cancer cell lines"

### 1 Supplementary Figures

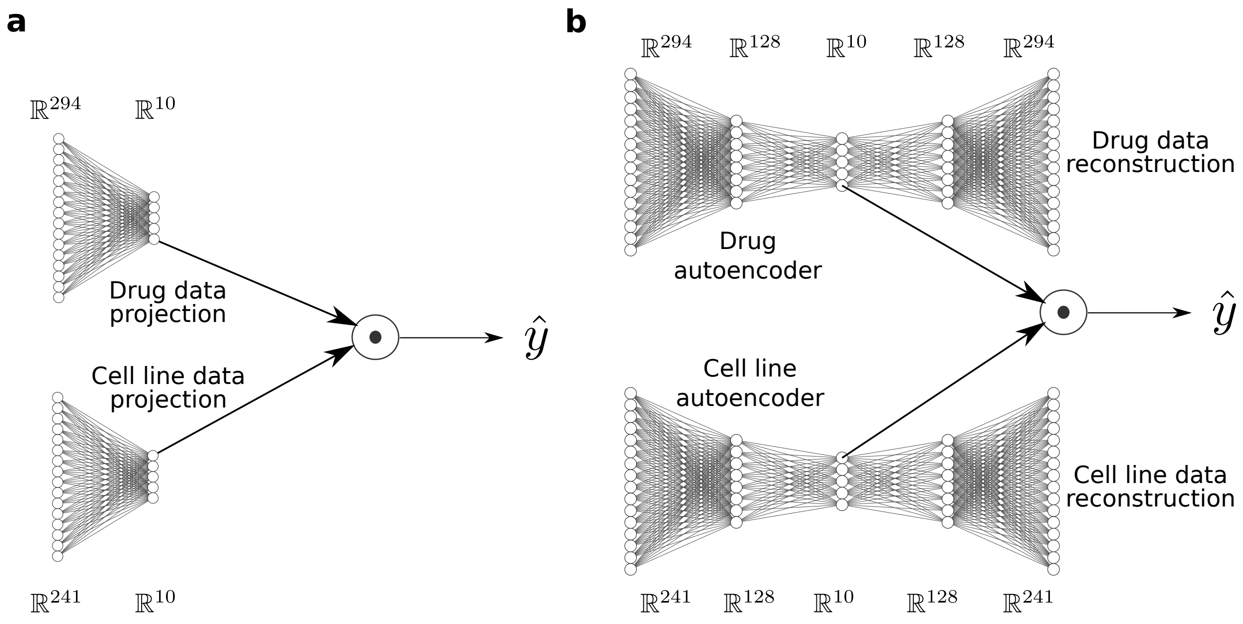


**Figure S1. Architecture of the models used for comparison. (a)** Linear matrix factorization model (Lin MF). First, drug and cell line data are linearly projected to 10-dimensional hidden representations. Prediction of the response of a cell line to a drug $\hat{y}$ is obtained by applying the dot product to the corresponding hidden representations. **(b)** Non-linear extension of the basic linear model (Autoen MF). The dimensionality reduction is performed via autoencoders with one hidden layer. Prediction of the response of a cell line to a drug is again obtained by taking the dot product of the corresponding, 10-dimensional hidden representations. Drug and cell line data reconstruction errors are also included in the optimization goal.

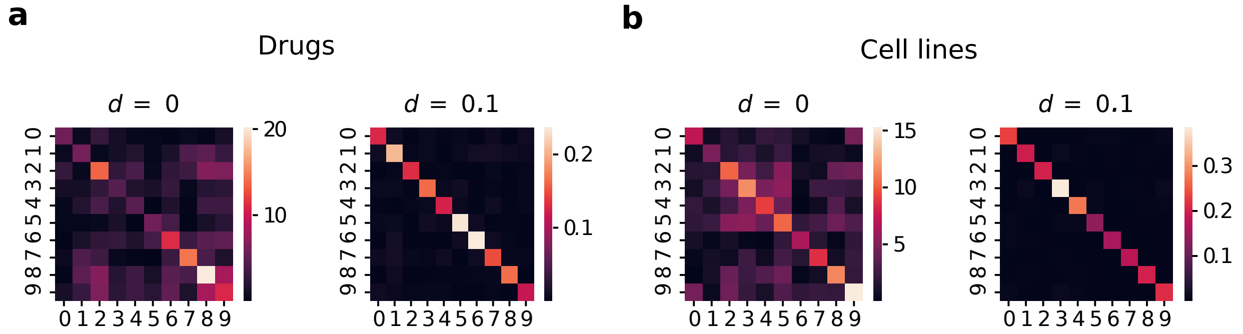


**Figure S2. Effect of dependence penalty $d$ shown for (a) drugs and (b) cell lines.** First, the drug and cell line data are passed to drug and cell line autoencoders of the DEERS model, respectively, obtaining 10-dimensional hidden representations of all drugs and cell lines used in the analysis. The covariance matrices are calculated across drugs and cell lines in the hidden space. The displayed covariance matrices correspond to the case where DEERS was trained without depencence penalty ($d = 0$), and with dependence penalty term $d = 0.1$.
